# Predictability of the community-function landscape in wine yeast ecosystems

**DOI:** 10.1101/2022.12.15.520418

**Authors:** Javier Ruiz, Miguel de Celis, Juan Diaz-Colunga, Jean CC Vila, Belen Benitez-Dominguez, Javier Vicente, Antonio Santos, Alvaro Sanchez, Ignacio Belda

**Author notes:** These authors contributed equally to this work (Javier Ruiz, Miguel de Celis).

## Abstract

Predictively linking taxonomic composition and quantitative ecosystem functions is a major aspiration in microbial ecology, which must be resolved if we wish to engineer microbial consortia. Here, we have addressed this open question for an ecological function of major biotechnological relevance: alcoholic fermentation in wine yeast communities. By exhaustively phenotyping an extensive collection of naturally occurring wine yeast strains, we find that most enologically-relevant traits exhibit a strong phylogenetic signal, indicating that the most relevant functions in wine yeast communities can be predicted from taxonomy. Further, we demonstrate that the quantitative contributions of individual wine yeast strains to the community function followed simple quantitative rules. These regularities can be integrated to quantitatively predict the function of newly assembled consortia. Besides addressing a fundamental open question in functional ecology, our results and methodologies provide a blueprint for rationally managing microbial processes of biotechnological relevance.

## Introduction

Microbes have been exploited for biotechnological purposes for millennia: from their original roles in food production to their myriad current uses in biomanufacturing. These microbial services are often delivered by complex and diverse ecological communities of microorganisms (Eng & Borenstein, 2019). Identifying the relationship between the taxonomic composition of microbial communities and the ecosystem functions they provide is critical if we wish to engineer microbial consortia and manage natural microbiomes (Dietze, 2018).

Ecological functions in complex communities emerge from individual contributions of each member species as well as from their ecological interactions. Thus, the first step to predictively linking taxonomy and ecosystem function is to determine whether the individual functional contributions of community members in isolation are in fact predictable from the phylogeny. This requires the existence of a strong phylogenetic signal in key functional traits, which is not always observed and, specially in microbes, cannot be taken for granted (Martiny et al., 2015). The second step is to learn how the functional contribution of each species will be altered by the presence of all other community members, i.e. to handle ecological interactions. Building predictive quantitative models of ecological interactions is a significant challenge (Friedman et al., 2017; Sánchez-Gorostiaga et al., 2019; Gowda et al., 2022), as interactions can have complex mechanistic origins and their number explodes with community size.

The critical goal of predictively linking taxonomic composition and ecosystem function leads us to two key questions that we must resolve: i) is there a phylogenetic signal in ecologically-relevant traits, which determine the individual contributions of a species to community functions? and ii) can we predict how the individual contributions of a species to a community function change in different community contexts, due to interactions with other community members?

As a model system to address these two questions, here we have studied wine fermentation -a system of profound socio-economic significance and one of the first microbiomes ever domesticated and exploited by humans-. In wine fermentations, the composition of the native yeast communities from grape musts (where a few dozens of yeast species can be found) changes over time as alcoholic fermentation proceeds. This ecological succession leads to the final dominance of *Saccharomyces cerevisiae*, which normally completes the consumption of the available fermentable sugars in grape musts (approx. 200g/L of glucose:fructose, in 1:1 ratio). Despite the complexity of this ecological succession by wine yeasts, wine fermentation is a reproducible process and it is therefore emerging as a model system for synthetic ecology studies (Boynton & Greig, 2016; Bagheri et al., 2020; Conacher et al., 2020).

To address the two questions outlined above in the wine fermentation system, here we first interrogate whether key functional traits of enological significance are predictable from the phylogeny. To this end we characterise the phylo-functional relationships among wine yeast species, measuring a total of 43 ecological and enological traits in a collection of 60 wine yeast strains belonging to 30 different species (spanning the expectable phylogenetic diversity of most commonly found wine yeasts). We find that the environmental preferences, the wine fermentation performance, and other key enological traits exhibit strong phylogenetic signals and can be imputed from a standard, fungal genetic marker (26S sequence). We then show that the contributions of individual wine yeasts to alcoholic fermentation can be predicted in different community contexts, as they follow simple quantitative rules that parallel the global epistasis concept in evolutionary genetics. Using methods that we have introduced in recent work (Diaz-Colunga et al., 2022) to infer the community-function landscape from a subset of observed consortia, we are able to quantitatively predict the wine fermentation function in complex wine yeast consortia, based on their community composition.

## Results

### A strong phylogenetic signal allows to predict relevant traits in wine yeasts

To address our first question -is there a phylogenetic signal for ecological and enological traits within the wine yeast community-it is necessary to assay a broad collection of yeasts covering the widest possible phylogenetic diversity within the wine yeast microbiome. Thus, we established a collection of 60 wine yeast strains (Table S1), including 30 different species (belonging to 22 genera and 10 families; both Ascomycota and Basidiomycota yeasts) (Figure 1A, Figure S1). All the yeast strains in our collection were isolated from wine environments, and it covers genera that are cosmopolitan (i.e. *Saccharomyces, Aureobasidium, Hanseniaspora, Metschnikowia or Lachancea*) as well as rare (*Cyberlindnera, Kluyveromyces*, or *Brettanomyces*) in wine fermentations (Figure S2). We then phenotyped this collection, focusing on traits that are relevant from an enological standpoint (Figure 1B). For instance, we measured the growth ability of the strains in a panel of 28 wine-related culture conditions (Table S2) to infer their environmental preferences, and we evaluated the fermentation performance of the strains by studying their contribution to the wine chemical profile (measuring 15 physical-chemical parameters, Table S3) of the resulting wines produced by each strain alone in laboratory scale fermentations. To our knowledge this represents the broadest functional catalogue of wine yeast species currently available (Table S4).

**Figure 1.**
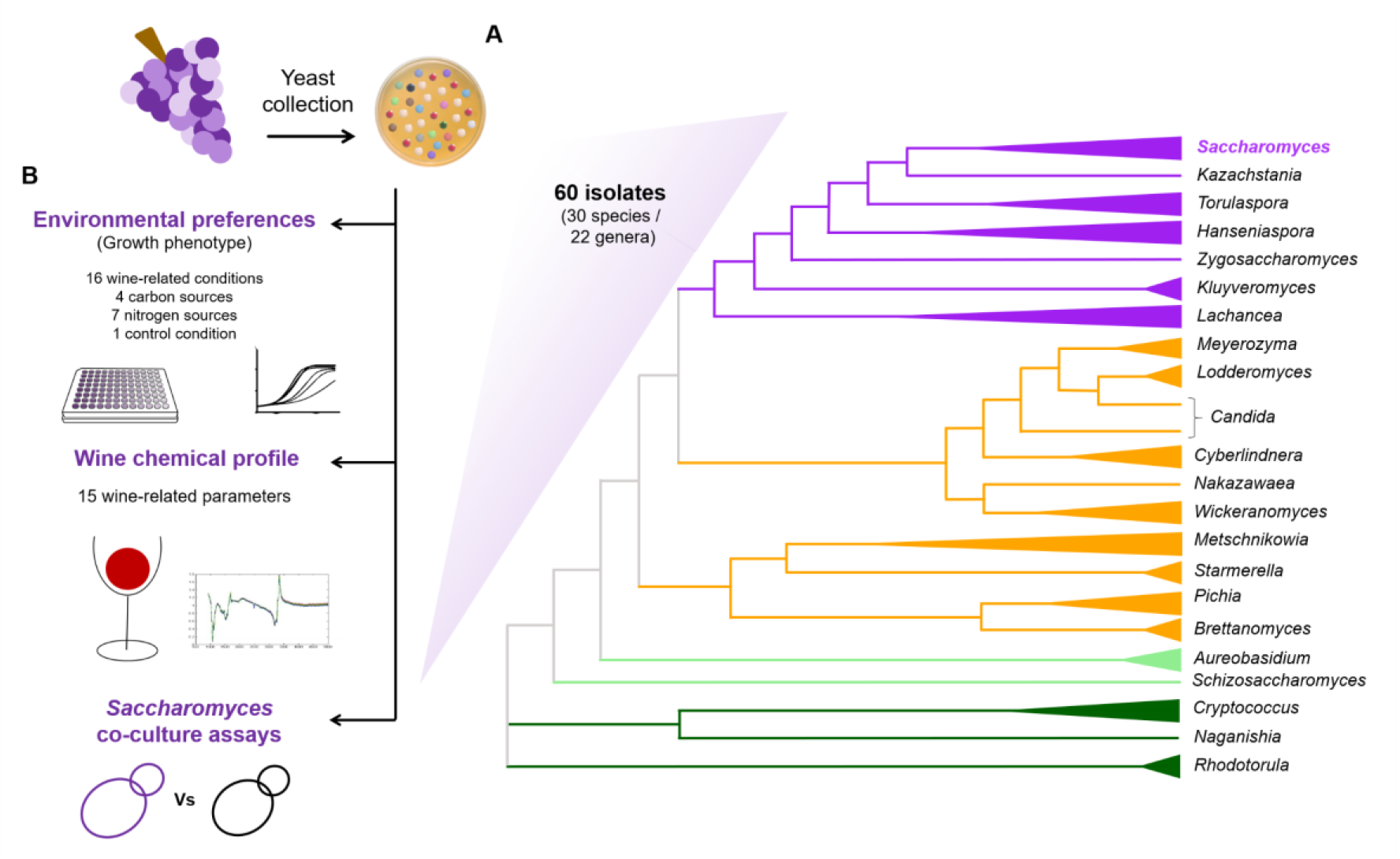
Composition and phenotyping strategy of the wine yeasts collection. **(A)** Schematic representation of the phylogenetic relationships of the 22 yeast genera included in the wine yeasts collection studied (based on the maximum likelihood phylogenetic tree presented in Figure S1, made using the partial 26S nucleotide sequence (D1/D2 domain of the 26S large subunit of rRNA gene) of the 60 strains of the study). Colours of the branches correspond to the lowest taxonomic level shared with *S. cerevisiae* (purple: same family, orange: same order, light green: same division, dark green: same kingdom). **(B)** We explored the most relevant enological traits of the 60 wine yeast strains included in our collection, by characterising: i) their environmental preferences, as their growth efficiency in a panel of 28 conditions (detailed in Table S2), including 16 wine fermentation conditions (modulating the sugar concentration, pH, antimicrobials, temperature, nitrogen or vitamins availability in Synthetic Grape Must (SGM)), 4 carbon sources (the three main organic acids found in the grape must, and glucose, as sole carbon sources) and 7 nitrogen sources (the six most abundant amino acids found in grape must, and ammonia, as sole nitrogen sources), and a synthetic medium as a growth control condition; ii) their fermentation performance, measuring 15 physical-chemical parameters in wines (detailed in Table S3) such as sugar consumption, ethanol production, organic acids production/consumption or total acidity produced, in laboratory-scale fermentations of SGM; and iii) their effect on the growth efficiency of co-cultures with *S. cerevisiae* (non-*Saccharomyces* x *S. cerevisiae*) in SGM.

This extensive functional characterization revealed a large phenotypic diversity across wine yeasts (Figure 2A-B). A first look at the heatmaps suggests a qualitative connection between the functional traits of the strains and their phylogenetic relationships, as we observe some distinguishable phenotypic patterns across taxonomic clusters both for environmental preferences (Figure 2A) and fermentation performance (Figure 2B). For instance, in wine fermentation conditions, Ascomycetes yeasts (purple and orange) exhibit, on average, higher growth efficiencies than basidiomycetes (light and dark green) (T-test, p=5.387e-07). Among Ascomycetes, Saccharomycetaceae strains (purple) have, on average, higher growth efficiencies than the non-Saccharomycetaceae strains (orange and light green) (T-test, p=1.604e-12), and, as expected at the species level, *S. cerevisiae* strains showed the highest growth efficiency (total variation in cellular density) in most fermentative conditions tested (Figure 2A), and it appeared as the only species able to deplete all sugars (glucose+fructose) in the laboratory-scale fermentations performed in SGM (Figure 2B).

**Figure 2.**
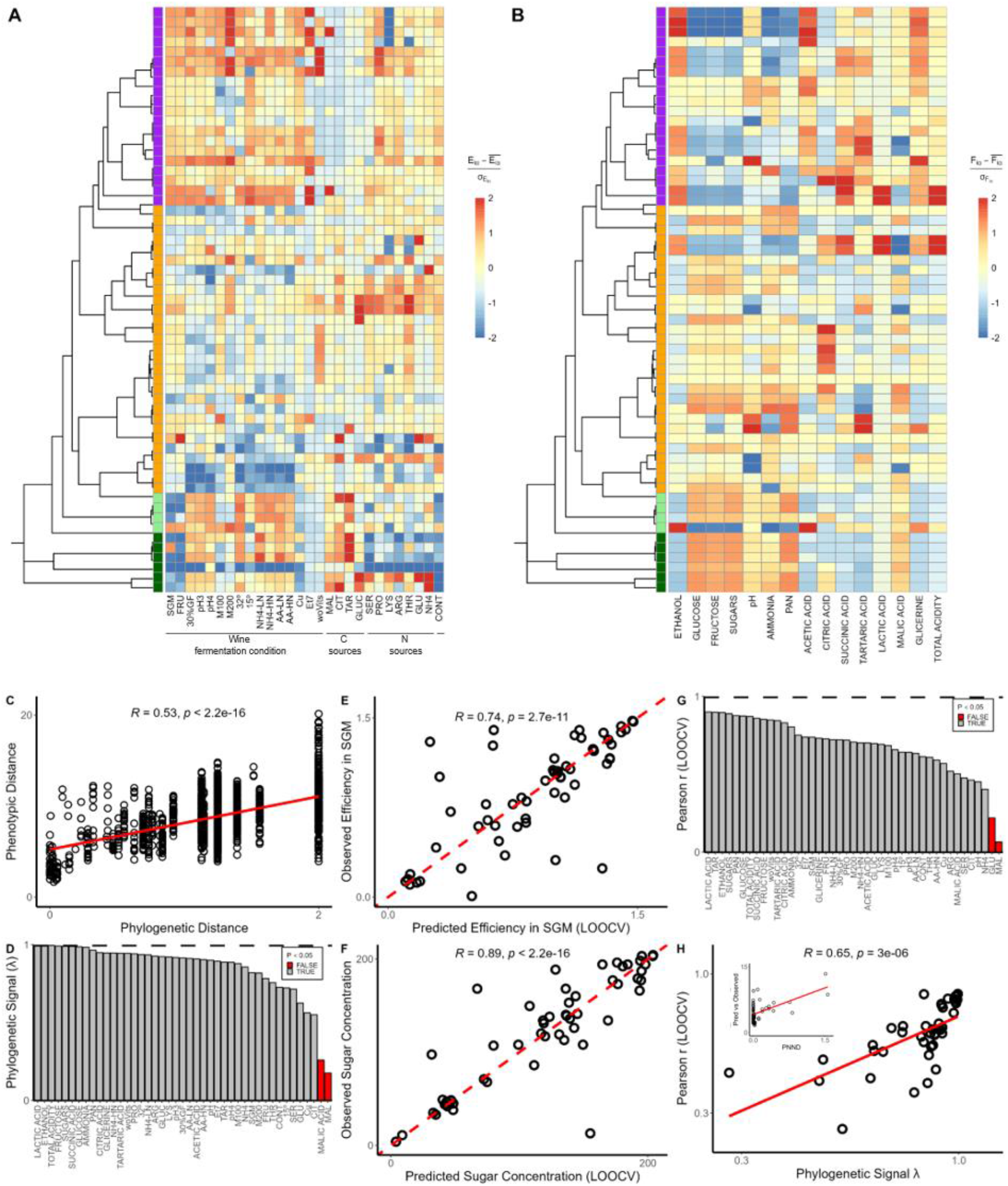
Environmental preferences and metabolite production of yeasts in wine fermentations can be predicted from phylogeny. Phylogenetic tree and trait heatmaps for 60 wine yeast strains showing **(A)** their environmental preferences, as growth efficiency in 28 different growing conditions (see Table S2 for detailed information about conditions tested), and **(B)** their wine fermentation performance, measuring 15 wine physical-chemical parameters after 168h of fermentation in Synthetic Grape Must (SGM) (see Table S3 for detailed information about the parameters measured). Table S4 contains the raw data used to plot the heatmaps. Trait values are centred and scaled for visualisation purposes. Colours to the left of the heatmap correspond to the lowest taxonomic level shared with *S. cerevisiae* (purple: same family, orange: same order, light green: same division, dark green: same kingdom. **(C)** We find a significant relationship between pairwise phylogenetic (cophenetic) distance and pairwise phenotypic distance (Euclidean distance in scaled trait values). **(D)** Phylogenetic signal (Pagel’s λ, hereafter λ) for all 43 traits measured (environmental preferences and fermentation performance). The λ parameter represents a continuous measure that varies from 0 to 1. λ = 0 indicates that the trait has evolved independently of the phylogeny whereas λ =1 indicates that the trait distribution amongst strains is highly influenced by their shared evolutionary history. 41/43 traits show a significant phylogenetic signal. **(E)** Predicted vs observed growth efficiency in SGM. **(F)** Predicted vs observed sugar (glucose+fructose) concentration after 168h of growth in SGM. All predictions (panels E and F) were made using phylogenetic imputation (Rphylopars) and tested using LOOCV. **(G)** Correlation coefficient between predicted and observed traits values for all 43 traits. A significant positive correlation is observed for 41/43 traits (Figure S4). **(H)** Traits with stronger phylogenetic signal are better predicted using phylogenetic imputation. In addition, phenotypic traits of strains whose nearest neighbour is further away (measured as phylogenetic nearest neighbour distance (PNND)) in the phylogenetic tend to be more poorly predicted (*R*=0.55; p=6.3e-6).

In order to quantify the relationship between phylogeny and phenotype, we next took advantage of the 26S sequence of the 60 yeast strains studied (Table S1). We inferred a maximum likelihood phylogenetic tree for our collection of 60 strains (Figure S1) and used this tree to quantify the phylogenetic (cophenetic) distance between every pair of yeast strain. We then determined the phenotypic similarity between every pair of strains, as the Euclidean distance between the scaled trait values, both from the environmental preferences and the fermentation performance of the studied strains. We find that there is a significant correlation (*R*=0.53, p=2.2e-16) between the phylogenetic distance (branch length) and phenotypic distance (Euclidean distance in trait space) between pairs of strains (Figure 2C, Figure S3). Furthermore, most phenotypic traits analysed in this work exhibited significant (lambda-test, p<0.05) phylogenetic signal (λ) (Figure 2D). Interestingly, we found that the growth efficiency in a synthetic medium with malic acid as the sole carbon source and the malic acid consumption during wine fermentations are the only two measured traits without a significant phylogenetic signal.

At this point, we wondered if the strong phylogenetic signal observed across wine yeast would allow us to quantitatively predict unobserved traits values in strains based on their phylogenetic position. To test this hypothesis, we leveraged R *Phylopars*, a tool for phylogenetic imputation of missing data (Goolsby et al., 2016). *Rphylopars* imputes phenotypic values in unobserved species by rerooting the phylogeny at the most recent common ancestor of the taxon with unobserved traits and the rest of the tree and then performing ancestral state reconstruction to obtain a maximum likelihood estimate for the trait value in this common ancestor (in this case under a Brownian model of trait evolution) (Bruggeman et al., 2009) The ability to predict unobserved trait values using phylogenetic imputation was determined by performing leave one out-cross validation (LOOCV) and then comparing observed trait values with those predicted by *Rphylopars* [see methods for a more detailed information]. In Figure 2E-2F we show a significant correlation between predicted and observed values for two different traits, growth efficiency (*R*=0.74, p=2.7e-11) and sugar consumption in SGM (*R*=0.89, p<2.2e-16). Similar correlations are observed for most of the other 41 traits measured in this study (Figure S4), with all but 2 showing significant Pearson correlation coefficients between predicted and observed trait values (Figure 2G). As expected, and consistent with simulation studies (Goberna & Verdu, 2016), in our empirical dataset, traits with a stronger phylogenetic signal (e.g., total acidity and ethanol production in SGM fermentations) showed a better predictability using phylogenetic imputation (Figure 2H). Predictions are also better at shorter phylogenetic distances, when there are closer relatives present in the tree (low phylogenetic nearest neighbour distance, PNND) that have already been characterised (inset in Figure 2H).

To check the accuracy of phylogenetically-based, functional trait predictions out of sample, we phenotyped 53 new strains (Table S5); and measured the growth efficiency parameter in SGM (Table S6) (the original 60 strain library was phenotyped again as a control). Using the phylogenetic imputation method trained with the original dataset of our 60 strains collection, here we show that it is possible to predict the growth phenotype of new strains, never tested before, with a similar accuracy as replicate measurements of the training set of strains in independent experiments (Figure S5).

### Mapping the community-function landscape allows to predict the function of complex yeast communities

*S. cerevisiae* is the dominant yeast in wine fermentations and it is able to complete the alcoholic fermentation process, depleting all fermentable sugars of grape musts on its own. However, its ecological partners (non-*Saccharomyces* species) could also play an important role in wine ecosystem functioning, either through direct contributions to the fermentation process, through interactions with *S. cerevisiae*, or both. Importantly, the contribution of each of these yeasts to the wine fermentation function is likely to depend on interactions with the other community members. Mapping out these interactions is essential if we wish to quantitatively predict how the composition of the yeast community affects the function of the wine ecosystem. To explore this question, we would have to place these wine yeasts in different community contexts, and determine how their contribution to the sugar consumption (as the main function of the wine ecosystem) varies as the composition of the community changes.

The number of potential background communities one may form out of our 60-strain library exceeds 10^18^. To be able to tackle this problem empirically, we must therefore first reduce the dimensionality of the problem. To do this in an unbiased manner we selected a subset of 10 non-*Saccharomyces* strains from our collection based on their effect on the growth efficiency in co-cultures with *S. cerevisiae* (Figure S6), including 5 strains that lower the growth efficiency of co-cultures with *S. cerevisiae* (*Hanseniaspora opuntiae* Hop_1 (Hop), *Kazachstania unispora* Ku_1 (Ku), *Metschnikowia pulcherrima* Mp_3 (Mp), *Pichia kudriavzevii* Pk_3 (Pk), and *Wickerhamomyces anomalus* Wa_4 (Wa)); and 5 strains that do not reduce the growth efficiency of co-cultures with *S. cerevisiae* (*Aureobasidium pullulans* Ap_1 (Ap), *Lachancea thermotolerans* Lt_2 (Lt), *Schizosaccharomyces pombe* Sp_2 (Sp), *Torulaspora delbrueckii* Td_8 (Td), and *Zygosaccharomyces bailii* Zb_1 (Zb)). Then, we assembled a total of 176 background communities by random combinations of these 10 strains in communities of 2-6 species (with the only constraint of ensuring a similar prevalence of all the species in the set of communities assembled). We assayed the function (final fraction of sugars consumed) of these 176 background communities, both by themselves and by adding, in separate experiments, two *S. cerevisiae* wine strains (Sc_5 and Sc_8) (Figure 3A; Figure S13).

**Figure 3.**
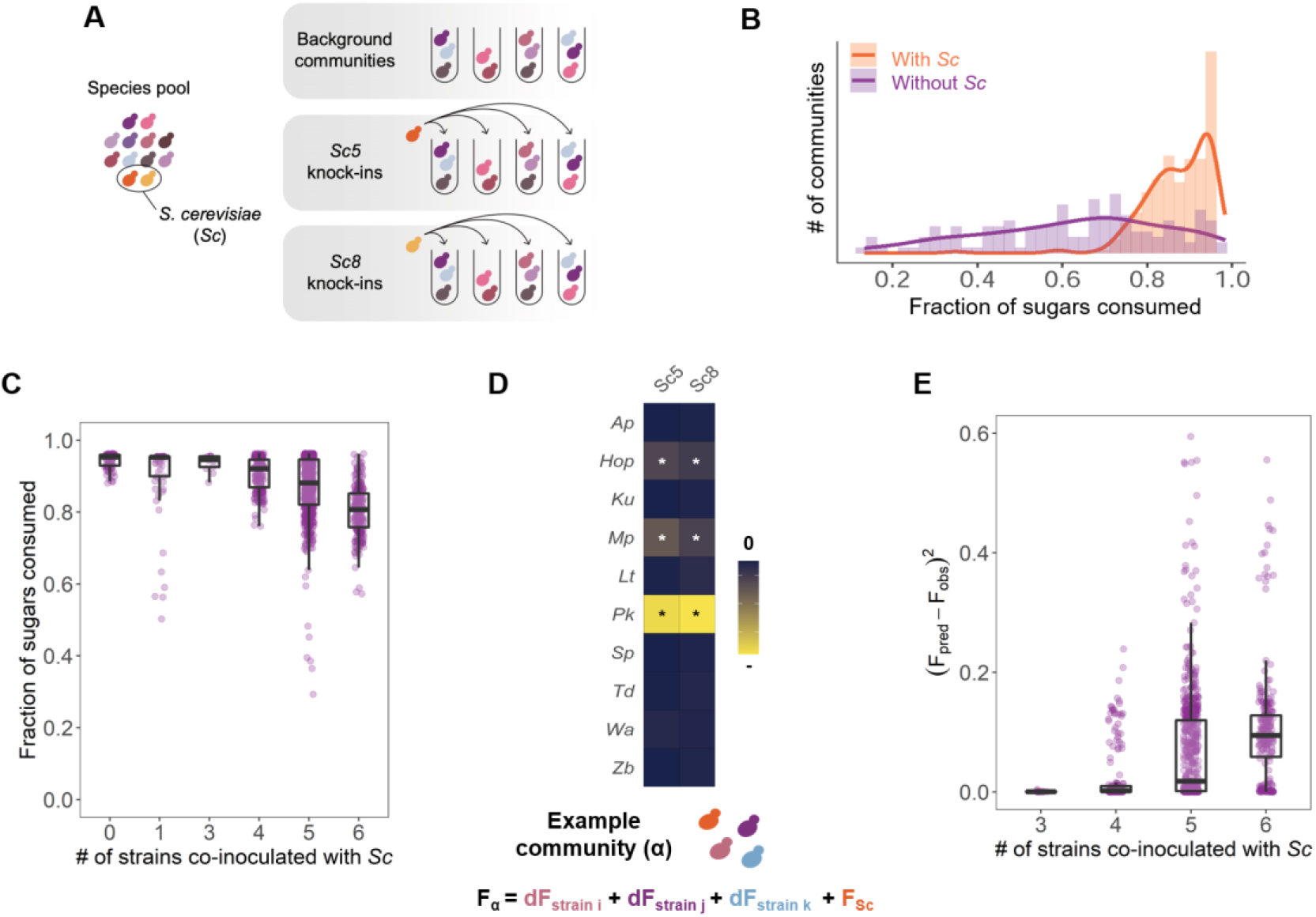
Ecological interactions affect the function and predictability of wine yeast communities. **(A)** To better understand how the complexity of a wine yeast community affects the fermentation performance of *S. cerevisiae*, we constructed synthetic communities of growing diversity. Based on the effect of the different species on the growth efficiency of co-cultures with *S. cerevisiae* (Figure S6), we have selected ten yeast strains (*Hop_1, Ku_1, Mp_3, Pk_3, Wa_5, Ap_1, Lt_2, Sp_2, Td_8, Zb_1*) to randomly assemble 176 communities (from 2 to 6 members) -named as background communities. These background communities were assayed by themselves and co-inoculated, in separate experiments, with two different *S. cerevisiae* strains (Sc5 and Sc8), constituting a total of 528 synthetic communities assayed. In addition, we assayed the function of all the 12 yeasts studied (ten non-*Saccharomyces* and the two *S. cerevisiae* strains) individually and in pairwise combinations. Thus, we performed a total of 606 assays, with three replicates, and after 168h of fermentation in SGM, we measured the fraction of sugars consumed (raw data on sugar consumption of each assay is available in Table S8). A schematic representation of the methodology followed to construct the communities can be found in the methods section and in Figure S13. **(B)** Despite the ability of *S. cerevisiae* strains to fully deplete sugars as a monoculture, we observe that certain background communities prevent *S. cerevisiae* to complete the consumption of fermentable sugars. **(C)** We find that, in our experimental conditions (since we pre-selected yeast strains based on their effect on *S. cerevisiae* growth efficiency), increasing the richness of the background community increases our chances to detect negative interactions in the community. Each dot represents the fraction of sugars consumed of each community in which *S. cerevisiae* strains were inoculated. We discard a direct effect of richness on community function, since Figure S8 shows no effect in the function of communities of growing richness, when they only contain the 5 strains that were pre-selected with a neutral or positive effect on the growth efficiency of co-cultures with *S. cerevisiae*. **(D)** Based on the individual functional effect (dF) of the strains on *S. cerevisiae* sugar consumption capacity, we established a simple additive model where the expected function of a given community was calculated by adding the decreasing effect on function caused by every strain included in that community. **(E)** The prediction accuracy of this model (calculated as the squared absolute difference between the observed and the predicted function value of the function) decreases as the diversity of the communities increases, failing to predict the ecological function of complex communities (Root Mean Square Error of each community complexity is: *RMSE*_*3*_=0.026; *RMSE*_*4*_=0.159; *RMSE*_*5*_=0.262; *RMSE*_*6*_=0.323).

As expected, communities containing *S. cerevisiae* consumed, on average, a higher fraction of sugars than the respective background communities which contained only non-*Saccharomyces* strains. However, some communities containing *S. cerevisiae* fell far from completing the fermentation, leaving, in some cases, ∼30% of residual sugars unconsumed (Figure 3B). We observe that, in our experimental conditions (since we pre-selected yeast strains based on their effect on *S. cerevisiae* growth efficiency), increasing the richness of the background community increases our chances to detect negative interactions in the community, preventing *S. cerevisiae* to complete the consumption of fermentable sugars (Figure 3C). Thus, we confirmed that the diversity of the background community where *S. cerevisiae* is added, play an important role in wine fermentation performance.

A key question is then to what extent interactions among community members play a role in the quantitative function of these communities, or if, alternatively, knowing the individual effect of yeast strains on the sugar consumption capacity of *S. cerevisiae* (Figure 3D) can be enough to predict the function of complex communities. We observed that the presence of yeast strains that caused a significant negative effect on *S. cerevisiae* sugar consumption capacity (Hop, Mp and Pk) can also anticipate a loss of function in complex communities, since they are more frequently found in communities where *S. cerevisiae* is not able to complete the consumption of sugars in SGM fermentations (Figure S7). Thus, knowing the individual effect of yeast strains can be useful to qualitatively infer their effect on the function of complex communities. However, exploring the simplest possible model -where the function of the community is calculated by the sum of the neutral or detrimental effect (d*F*) of each strain in the sugar consumption capacity of co-cultures with *S. cerevisiae* (Figure 3D)-we are far from quantitatively predicting the function of complex communities (three, four, five or six strains co-inoculated with *S. cerevisiae*) (Figure 3E). This suggests the existence of a complex network of interactions -beyond the individual interaction of yeast strains with *S. cerevisiae*-that ultimately shape the function of wine yeast communities, and that should be considered to accurately predict their function.

While interactions among yeasts can be important for community function, their number rapidly explodes with community size, and it is not obvious how one should integrate them even if we knew them. A recent work by Diaz-Colunga and colleagues (2022) has found that the collective effect of these interactions is the emergence of simple linear models that predict how each species affects ecosystem function when placed in different community contexts. Moreover, the function of a complex microbial community can be predicted by integrating these linear models. These linear models relate the change in ecosystem function caused by the inoculation of a new focal strain into a new background community (*functional effect*, d*F*), and the function of a background community (background community function, F). We sought to apply this theoretical framework to address the challenge of predicting the functional effect of different wine yeast strains in the presence of interactions. Our experimental approach allowed us to compare the fraction of sugars consumed by a background community with the change in that fraction produced by both the two *S. cerevisiae* strains (Figure 4A), and by all the 10 non-*Saccharomyces* strains assayed (Figure 4B). In this way, we can characterise the ecological effect of these strains by analysing the slope and intercept of the linear models that predict their functional effects in different community backgrounds (these models are referred to as Functional Effect Equations, or FEEs). As expected, we observe high intercepts for both fits of *S. cerevisiae* strains, since they showed the highest individual function (they are able to mostly deplete the 100% of sugars in SGM in isolation). However, note that if *S. cerevisiae* were able to consume all sugars regardless of the composition of its accompanying community, its functional effect equation would be well described by a line of the form *y* = 1 - *x* (dashed lines in Figure 4A). Instead, we found that the functional effect equations of both *Sc_5* and *Sc_8* fall below this line, reflecting that *S. cerevisiae* strains are unable to complete wine fermentations in some ecological contexts.

**Figure 4.**
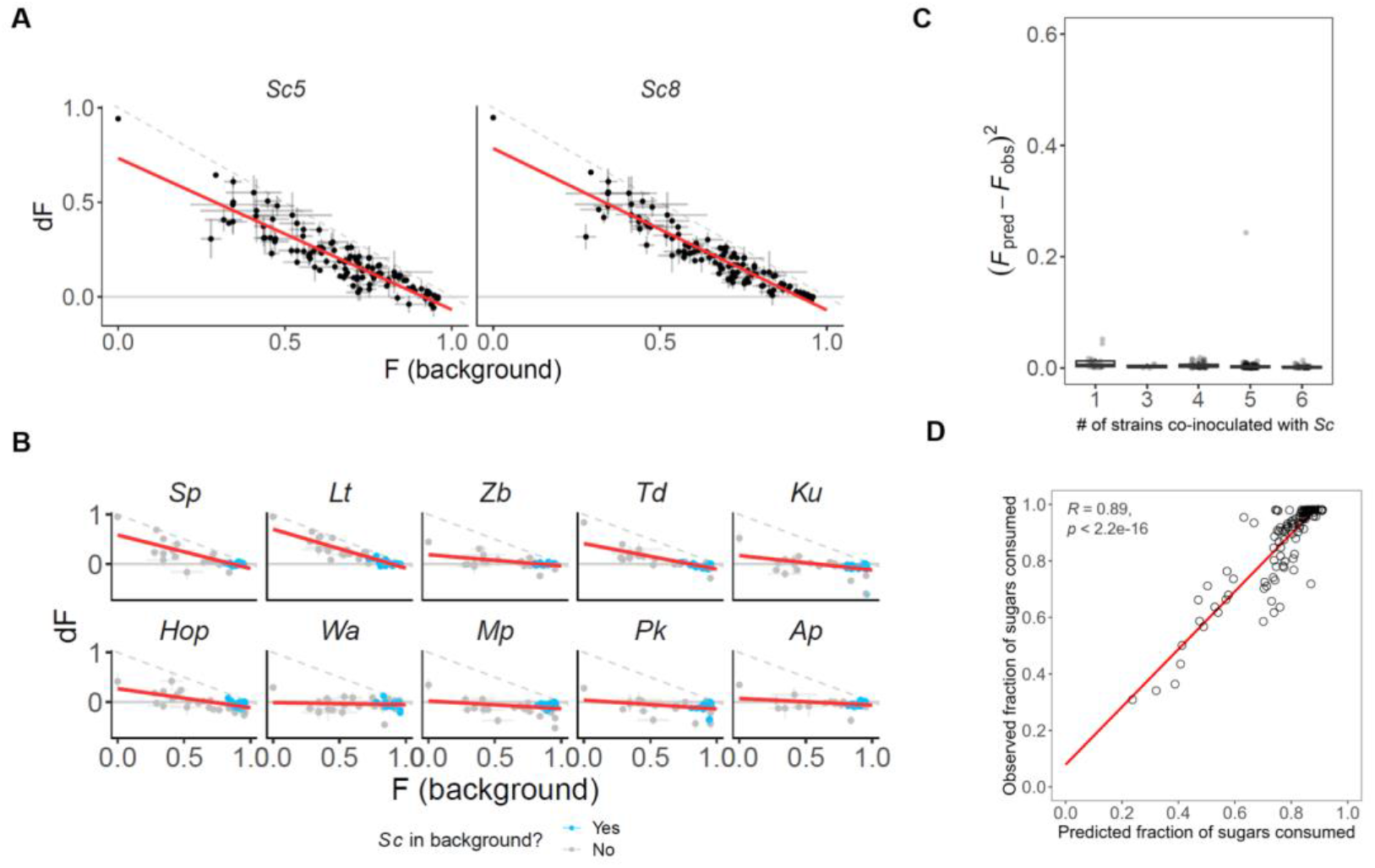
By mapping the ecological interaction occurring within complex wine yeast communities we can quantitatively predict their function. The contribution of the different yeast strains to wine ecosystem function depends on both their own sugar consumption capacity and the function of the background community to which it is added. **(A)** We show the functional effect (dF) of the two *S. cerevisiae* strains (Sc_5 and Sc_8) by representing the change in community function caused by the inoculation of *S. cerevisiae* strain in various backgrounds. The functional effect is defined as the difference in the fraction of sugars consumed by the community with *S. cerevisiae* and the fraction of sugars consumed by the same community without the *S. cerevisiae* strain (F(background)). Dots and error bars represent means and standard deviations of the three replicates. Dashed lines represent the functional effect of the strains if its inoculation would cause the full completion of sugar consumption (described as *y = 1 - x*). We refer to the linear fits as the FEEs. **(B)** FEEs of the ten non-*Saccharomyces* strains assayed in this experiment. Consistent with similar findings in a wide range of other ecosystems (Diaz-Colunga et al 2022) each of the non-*Saccharomyces* strains has its own FEE. Blue dots represent the background communities that do contain *S. cerevisiae*. Thus, we can study the effect on *S. cerevisiae* fermentation performance on its ecological context (Figure S9). Following the methodology described in Diaz-Colunga et al., (2022), we can predict the function (final fraction of sugars consumed) of this community by sequentially adding the functional effects of each strain that composes each community. **(C)** This method allows us to accurately predict the function of the community regardless of its complexity. **(D)** In order to test the prediction accuracy of this FEEs-based model to predict the function of any randomly assembled community, we assembled 131 new communities (never assayed before) and measured the fraction of sugars consumed after the fermentation. We compared the predicted function of these communities with the measured function, finding a high correlation between the observation and the prediction (*R*=0.89) (Table S9).

Likewise, Figure 4B shows that the functional effects of the ten non-*Saccharomyces* strains assayed are also well described by linear fits. The high fermentative yeasts *S. pombe* (Sp), *L. thermotolerans* (Lt) and *T. delbrueckii* (Td) exhibit a FEE that is very similar to that of the two *S. cerevisiae* strains. However, some strains (mainly Mp and Pk, but also Wa and Ap) exhibited a marked negative effect on the function of the community, and the function of the background community has little to no effect in magnitude of their impact. Other strains, such as Hop -or more subtly Ku*-* have functional effects that are positive in low-functioning backgrounds but negative in high-functioning backgrounds (i.e. communities with *S. cerevisiae* strains).

This approach was useful to more accurately characterise the qualitative contribution of each strain in different ecological contexts, allowing us to estimate if they might facilitate or prevent *S. cerevisiae* from completing fermentation. But further, here we demonstrate that by concatenating the FEEs of each strain member of the community -as described by Diaz-Colunga et al. (2022)-, we can quantitatively predict the function of any randomly assembled wine yeast community, independently of their diversity level (Figure 4C). To double check the robustness and accuracy of this model, we assembled and assayed 131 new yeast consortia that have not been assayed in the first experiment used to fit the model, comparing the predicted and observed values for their sugar consumption capacity (Table S9). This assay was conducted under identical conditions than the first experiment but it was carried out by a different member of the laboratory. As shown in Figure 4D, we found a significant strong correlation between predicted and observed functions (*R*=0.89). Thus, we confirm that knowing the individual behaviour of each species in different ecological contexts, we can quantitatively link the taxa composition of complex communities to the ultimate function they provide.

## Discussion

Using wine fermentations as a model ecosystem, we have addressed a major question in microbial ecology and evolution, studying how taxonomy shapes ecological function. We first show that key functional traits of enological significance exhibit strong phylogenetic signal. The phylogenetic conservatism of microbial traits is required if we wish to predict the presence of different ecological functions in a community from its taxonomic composition. Yet, this cannot be taken for granted (Martiny et al., 2013): the genetic and physiological plasticity of microorganisms and their rapid evolutionary adaptation rates could suppose a weaker phylogenetic signal in particular traits or systems (Blomberg et al., 2003; Hertz et al., 2013; Krause et al., 2014). Our results indicate a strong correlation between the phylogenetic and phenotypic distance of wine yeast species (Figure 2C; Figure S3), detecting a significant phylogenetic signal for almost all traits analysed (Figure 2D). Martiny et al. (2013) discussed that complex traits are more phylogenetically conserved than simpler traits that involve one or few genes, such as the consumption of a specific nutrient. In our work, most traits studied are based on the ability of yeasts to grow in synthetic grape must, i.e. fermentation capacity. This capacity is associated to some specific events in the evolutionary history of yeast (i.e. Whole Genome Duplication and the loss of the Respiratory Complex I) (Dashko et al., 2014), providing a plausible explanation for why this trait is strongly conserved in the phylogeny. Interestingly, the only two traits analysed that did not show a significant phylogenetic signal are related to malic acid consumption capacity of yeasts. The use malic acid as carbon source requires the yeast to import and metabolise L-malate, using a malate transporter (*S. cerevisiae* lacks an active transport system for L-malate), and a decarboxylating malate dehydrogenase (the malic enzyme), that can be mitochondrial (less active) or cytosolic. More studies will be necessary to explore the evolutionary origin of malic acid consumption capacity in phylogenetically distant wine yeast species (Saayman & Viljoen-Bloom, 2006).

Additionally, we demonstrate that phylogeny-based imputation (Goberna & Verdú, 2016), can be a valuable tool for yeast researchers, as the strong phylogenetic signal observed allowed us to accurately predict both the environmental preferences and the enological properties of a broad collection of yeast species. During the last decade, high-throughput sequencing studies in wine fermentations revealed a greater than expected yeast diversity in wine fermentations (Bokulich et al., 2014; Knight et al., 2015; Liu et al., 2020; de Celis et al., 2022), increasing the number of potentially relevant species to be characterised. Thus, we stand out the relevance of the dataset and predictive tools provided in this work to easily predict wine-relevant traits of new taxa, only requiring their phylogenetic (26S sequence) information.

Several previous works have explored in detail the intra-specific phenotypic diversity of some relevant wine yeasts, such as *S. cerevisiae* (Camarasa et al., 2011), *L. thermotolerans* (Hranilovic et al., 2018) or *T. delbrueckii* (Silva-Sousa et al., 2022). Here, looking at the inter-species diversity, we provide information about ecologically and enologically-relevant traits of 60 wine strains belonging to 30 different yeast species. We hope that our yeast collection and its phylo-functional characterization would also be a useful resource for wine researchers to guide future wine yeast selection programs. In this regard, we highlight some outstanding results, that may help, for instance, to contribute to mitigate the effects of climate change in the wine industry, such as the capacity of *T. delbrueckii* to grow in vitamins-deficient grape musts, or to modulate wine acidity by producing lactic acid (*L. elongisporus* and *L. thermotolerans*) or degrading malic (*H. osmophila, L. elongisporus* and *P. kudriavzevii*) or tartaric acid (*C. amylolentus*).

Once we have established that there exists a strong link between taxonomy and function within the wine yeast microbiome, we sought to understand the effect of inter-species interaction patterns in modulating the functional contributions of each community member, and in shaping the ecological function of complex wine yeast communities. *S. cerevisiae* becomes the dominant yeast during wine fermentations, as it is able to rapidly complete wine fermentations, consuming all fermentable sugars from the grape must by its own. This ability is result from the adaptation of *S. cerevisiae* to the challenging conditions of wine fermentations (Marsit & Dequin, 2015; García-Ríos & Guillamón, 2019), due to a number of genomic features that wine strains acquired over their domestication process (Belda et al., 2019). However, we observed that some combinations of yeast strains caused a negative effect on *S. cerevisiae* function (Figure 4B; Figure S7). This suggests that the accompanying yeast species -despite their limited sugar consumption capacity-, can potentially prevent *S. cerevisiae* from completing wine fermentation. The effect of these species on the community function cannot be accurately explained by the additive effect of the individual interaction patterns of each accompanying species with *S. cerevisiae* (Figure 4E). This indicates the importance of interactions in determining the effect of complex microbial communities on *S. cerevisiae* fermentation performance. Supporting this idea, a recent work of Conacher et al. (2022) has evidenced the presence of high-order interaction within the wine microbiome, demonstrating that the inoculation of *S. cerevisiae* with complex yeast communities disclose transcriptional responses not explained by the transcriptional responses observed in pairwise co-culture.

If we aspire to engineer microbiomes for biotechnological purposes, we need to be able to predict the function of complex communities, guiding the search for the best possible combination of species. In principle, quantitatively predicting the fraction of sugars consumed by a particular combination of yeast species during wine fermentations, might seem like a challenging task, requiring extensive characterization of the chemical and physiological mechanisms through which yeast strains may interact with one another. However, following the methodology presented by Diaz-Colunga et al., (2022), we demonstrated that the contribution of different wine yeast species to the function of complex communities can be described by linear patterns based on their own sugar consumption capacity in different ecological contexts. We show that, by characterising these patterns we could not only improve our understanding of the ecology of wine yeasts (Figure 4A-B; Figure S10), but to use this information (FEEs) to fit a model that allows us to quantitatively predict the ecological function of the wine yeast communities. Herein we have taken advantage of the theoretical framework established by Díaz-Colunga et al. (2022) by applying the described methodology to improve the understanding of a highly valuable biotechnological process such as wine fermentation.

Additionally, given the success we had in using phylogenetic imputation to predict ecologically relevant traits in wine yeasts (Figure S4), we asked whether the FEE slope and intercept could also be interpreted as complex ecological traits, and thus, can be predicted in the same manner. Figure S11 shows a marginally significant correlation between observed and predicted FEEs parameters (*R*=0.54, for both slope and intercept), but it seems that the functional effect of yeast strains in complex communities cannot be clearly predicted by phylogeny. The more limited size (n=12) of our strains collection assayed in complex consortia, and the strong impact of the function of the background communities might explain this lack of predictability on the ecological effect of the yeast strains. Moreover, we also looked for individual traits (among those described in Figure 2) that can be somehow related with the ecological effect of strains (slope and intercept in the FEEs) in wine yeast communities. Although further work will be necessary to further establish causal links between the metabolic traits of a given species and its contribution to the ecosystem function, we found that the ethanol production capacity, the consumption of organic nitrogen and the glucose:fructose consumption ratio, are important predictors of the effect that one strain may have to the function of wine yeast communities (Figure S12). In particular, we will address future works to elucidate the molecular and ecological mechanisms of action of those yeasts found here as leading to stuck wine fermentations (mainly *P. kudriavzevii, H. opuntiae* and *M. pulcherrima*), but, by now, we finish this work providing evidences that, even in microbial communities usually dominated by a single highly-adapted species, interspecies interactions play a crucial role in ecosystem function.

To conclude, with this work we want to encourage the use of fermented foods as model systems (Wolfe & Dutton, 2015) to address fundamental questions in ecology and evolution, but also to adopt the conceptual frameworks of these disciplines for a more integrated understanding of microbial-based industrial processes.

## Methods

### Yeast strains and molecular identification

Yeast strains used in this study are listed in Table S1 (main collection of 60 yeast strains used in the study) and Table S5 (list of 53 additional yeast strains used for double checking phylogeny-based phenotypic predictions (Figure S5)). All yeast isolates were identified by sequencing the D1/D2 domain of the 26S large subunit of rRNA gene, with the forward NL-1 primer (5′-GCATATCAATAAGCGGAGGAAAAG-3′) and the reverse NL-4 (5′-GGTCCGTGTTTCAAGACGG-3′) primer. After Sanger sequencing, the sequences obtained were compared and identified by BLAST-search and submitted to the GenBank database (NCBI accession numbers are detailed in Table S1 and Table S5). *Sabouraud* medium (Oxoid, Hampshire, UK) was routinely used for handling of the strains.

### Estimation of population prevalence and relative abundance of yeast genera in wine samples

The population prevalence and relative abundance patterns of the yeast genera represented in our yeast collection (showed in Figure S2) were calculated using the dataset reported in de Celis et al. (2022), where we analysed a total of 144 wine fermentation samples following an ITS-amplicon sequencing strategy. Raw sequences (fastQ files) are available at NCBI (Bioproject: PRJNA814622); and, details about DNA extraction, sequencing, bioinformatics and taxonomic assignment are described in the work by de Celis et al. (2022). In the particular case of the genus *Metschnikowia*, its population prevalence and relative abundance figures were obtained from the work of Vicente et al. (2020).

### Phylogenetic Tree Reconstruction

Sequences from the D1/D2 domain of the 26S large subunit of rRNA gene of the 60 yeast strains (NCBI accession codes detailed in Table S1) were used to construct a phylogenetic tree (Figure S1). We aligned the 60 sequences using the *msa* package (v3.16) (Bodenhofer et al., 2015) in R (v 4.2.2), implementing the ClustalW algorithm. The alignment was trimmed using trimAl (automated1) to remove poorly aligned regions (Capella-Gutiérrez et al., 2009). After trimming we inferred a maximum likelihood phylogenetic tree using IQTREE 2 with 1000 bootstrap replicates (‘iqtree2 -s alignment.phy -bb 1000 -redo’). The consensus tree was rooted using the Basidiomycota clade as an outgroup as this represents the deepest known taxonomic division amongst all yeast strains. An ultrametric tree was then obtained using the chronos function in the *ape* R package (v5.6-2) and all subsequent analysis was performed on this tree. Heatmaps showing trait distributions across the phylogenies (Figure 2A-B) were constructed using the *ggtree* R package (v3.16) (Yu, 2020).

### Study of environmental preferences (growth assay) of the yeast collection

To study the environmental preferences of our yeast collection (represented in Figure 2A), the 60 yeast strains were assayed in a panel of 28 growing conditions of enological interest (detailed information about the composition of culture media and growing conditions is available in Table S2), in brief: 16 wine fermentation conditions (growth assays in synthetic grape must-based media), the capacity to grow using the most relevant organic acids of grape musts as sole carbon sources (malic acid, citric acid, and tartaric acid; plus glucose as a control), the most abundant amino acids of grape musts as sole nitrogen sources (lysine, serine, threonine, arginine, glutamic acid, proline; plus ammonia as a control), and a minimal medium (Yeast Nitrogen Base (YNB, BD Difco™, USA)) without amino acids and (NH_4_)_2_SO_4_ (1.7 g/L); glucose (200 g/L); (NH_4_)_2_HPO_4_ (0.24 g/L); amino acids 20X stock solution (10.79 ml)) as an standard growth control condition.

Synthetic Grape Must (SGM) was prepared as described by Henschke & Jiranek (1993), with the modifications detailed below. Firstly, vitamins (20x), amino acids (20x), trace elements (1000x) and anaerobic factor (200x; ergosterol, Tween 80 and ethanol) stock solutions were prepared separately, sterilised by 0.45 μm filtration and stored at 4 ºC (maintaining the proportion of the original recipe). To prepare the SGM, the stocks solutions (50 mL of vitamins; 10.79 mL of amino acids stock; 1 mL of trace elements stock and 5 mL of anaerobic factor stock) were dissolved in H_2_O followed by glucose (100 g/L), fructose (100 g/L), malic acid (3 g/L), potassium L-tartrate (2.5 g/L), citric acid (0.2 g/L), KH_2_PO_4_ (1.14 g/L), MgSO_4_·7H_2_0 (1.23 g/L), CaCl_2_·2H_2_O (0.44 g/L) and (NH_4_)_2_HPO_4_ (0.24 g/L). Then, pH was set to 3.5 with KOH and SGM was sterilised by 0.45 μm filtration. Yeast Assimilable Nitrogen (YAN) was adjusted to 200 mg N/L (60 mg N/L of ammonia-nitrogen ((NH_4_)_2_HPO_4_) and 140 mg N/L of amino acids-nitrogen). This SGM medium was used for assaying the growth capacity of yeasts in 16 wine fermentation conditions, with the required modifications in each case (Table S2). The growth capacity using different carbon sources were assayed in Synthetic Medium (SM) prepared with YNB (1.7 g/L) and 1g/L of the corresponding carbon source (malic acid, citric acid, tartaric acid, and glucose as a control). Likewise, the growth capacity using different nitrogen sources was assayed in SM prepared with Yeast Carbon Base (1.7 g/L) (YCB, BD Difco™, USA), and 200 mg N/L of the corresponding nitrogen source (serine, proline, lysine, arginine, threonine, glutamine, and ammonia as a control). All media were sterilised by 0.45 μm filtration.

To perform these assays, strains were streaked out on *Sabouraud* medium plates and grown at 28°C during 24h. Then, strains were pre-cultivated for 24 hours on 250 μL of SGM in a 96-well microplate. After that, the strains were inoculated, as biological triplicates, at a final optical density (OD _600nm_) of 0.01, in 96-well plates filled with 250 μL of the corresponding medium. Assays were performed at 25ºC (except the fermentation assays at low (15ºC) and high (32ºC) temperatures) under orbital shaking set at 100 rpm. Yeast growth was monitored by measuring OD _600nm_ at different time points during 72 hours, using the microplate reader Varioskan Flash Multimode Reader (Thermo Scientific, USA). Growth efficiency (total variation in cellular density) was extracted by using *GrowthRates* package (v0.8.4) in R (Hall et al. 2014), adjusting the growth curves to a Baranyi model.

### Synthetic grape must microvinification assays

The fermentation performance of the 60 yeast strains of our collection (that is: their contribution to the chemical profile of wine, determined by measuring 15 physical-chemical parameters at the end of fermentations) was studied in microvinification assays using SGM (results presented in Figure 2B). Firstly, strains were precultured during 24h in 12-well plates containing 3 mL of SGM at 25ºC under orbital shaking (100 rpm). Then, strains were inoculated in SGM at a final OD_600nm_ of 0.01. All fermentations were performed as biological triplicates at 25 ºC (100 rpm agitation) in 15 mL plastic tubes filled with 14 mL of SGM. After 168h of fermentation, cultures were centrifuged at 7,000 rpm for 10 minutes to remove biomass. Then, supernatants were stored at -20ºC until further analysis to determine their fermentation performance (Table S3; Table S4). Wine pH, total acidity, volatile acidity, and the concentration of ethanol, malic acid, lactic acid, glucose and fructose were determined by the near infrared spectroscopy, using an infrared analyser (Bachus 3 MultiSpec, *Tecnología Difusión Ibérica*, S.L, Spain). The concentration of ammonia and primary amino nitrogen (PAN) were determined enzymatically by using the appropriate kits (Biosystems, Barcelona, Spain) and a spectrophotometric autoanalyzer Y15 (Biosystems, Barcelona, Spain).

### Phylogenetic signal analysis

The phylogenetic (cophenetic) distance between yeast strains was calculated as the sum of branch lengths between pairs in the maximum likelihood tree (Figure S1). Phenotypic distance between yeast strains was calculated as the Euclidean distances on the scaled trait values, both from the environmental preferences and the fermentation performance of the studied strains (Table S4; shown in Figure 2A-B). For each trait we quantified the phylogenetic signal (Pagel’s λ) by using the *phylosig* function from the *phytools* R package (v0.1-9). Pagel’s λ parameter represents a continuous measure that varies from 0 to 1. λ = 0 indicates that the trait has evolved independently of the phylogeny whereas λ =1 indicates that the trait has evolved according to a Brownian motion model of evolution. λ significant values close to 1 indicate that the trait analysed shows phylogenetic signal.

### Predicting Wine Yeast Traits with Phylogeny

For each of the 43 phenotypic traits measured (28 growing conditions and 15 physical-chemical parameters), we tested whether it is possible to predict the traits of a novel strain in the phylogeny using the phylogenetic imputation algorithm that assumes a Brownian motion model of evolution. To do that we used the *phylopars* R package (Goolsby et al., 2016). Predictions were performed using the Leave One Out Cross-Validation (LOOCV) method. Specifically, we removed all traits values associated with a given strain before fitting and imputed the missing value using the phylopars algorithm. Observed and predicted traits values were then compared. The predictive accuracy of the algorithm was quantified for each trait by calculating the Pearsons’ correlation coefficient between predicted and observed values. A wider collection of yeast strains (the 60 strains already characterised, plus 53 new strains; Table S5) was used to check the accuracy of phylogenetically-based predictions. The growth efficiency parameter in SGM was measured for the 113 strains (Table S6). Thus, 60 strains have been assayed in two independent batches of experiments, and 53 strains have been assayed once. We used this information to compare the prediction based on the phylogenetic imputation method trained with the original dataset of 60 strains, and the accuracy of the replicate measurements of the same strain in independent experiments. This phylogenetic imputation algorithm was also used to discuss the predictability of the main FEEs parameters (slope and intercept) based on phylogeny, as shown in Figure S11.

### Co-culture assays of S. cerevisiae and non-Saccharomyces strains

We measured the effect of the non-*Saccharomyces* strains on the growth efficiency of co-cultures with *S. cerevisiae* (results shown in Figure S6; Table S7). Co-culture growth assays were performed by co-inoculating the 60 strains of the study with *S. cerevisiae* Sc_5 strain (1:1 proportion, at a final OD_600nm_ of 0.02) in SGM media. Sc_5 x Sc_5 co-culture was also assayed, as a control, at a final OD_600nm_ of 0.02. These assays were performed, in 6 biological replicates, as described previously for the environmental preferences growth phenotyping (in 96-well plates filled with 250 μl of SGM, at 25ºC with 100 rpm agitation, for 72 hours), and the growth efficiency (total variation in cells density) values were obtained from growth curves as previously described. The effect of each strain on the co-culture function (growth efficiency) was expressed as dF=F_i+Sc_ - F_Sc_, where F_i+Sc_ is the function of the co-culture when the strain i is co-inoculated with *S. cerevisiae* strain, and F_Sc_ is the function of *S. cerevisiae* single culture (Sc_5 x Sc_5 assay). This experiment allowed us to identify five strains that caused an increase on the co-culture function (Ap_1, Lt_2, Sp_2, Td_8, Zb_1) and five strains that caused a decrease on the co-culture function (Hop_2, Ku_1, Mp_3, Pk_3, Wa_4) compared to *S. cerevisiae* single culture. These strains were selected for the subsequent construction of synthetic yeast consortia, as explained below.

### Construction and study of synthetic consortia of wine yeasts

To test how the composition of a background community can affect the ecological function of *S. cerevisiae* in wine fermentation, we assembled a total of 176 synthetic consortia combining the ten non-*Saccharomyces* strains mentioned before (*Ap_1, Lt_2, Sp_2, Td_8, Zb_1, Hop_2, Ku_1, Mp_3, Pk_3, Wa_4*). The assembly of the synthetic consortia was carried out as represented in Figure S13: first we designed 16 background communities (between 3-6 species) by random combinations of the 10 non-*Saccharomyces* strains (with the only constraint of ensuring a similar prevalence of all the species in the set of communities assembled), then, we also added each of the 10 species on each background community, to determine their functional effect in different ecological contexts (the composition and function of all synthetic yeast consortia assayed can be found in Table S8). These 176 communities were then assayed by measuring their sugar consumption capacity in SGM (explained in the following section), both by themselves and by adding, in separate experiments, two *S. cerevisiae* wine strains (Sc_5 and Sc_8), resulting in a total of 528 combinations assayed as biological triplicates. All yeast strains (10 non-*Saccharomyces* and 2 *S. cerevisiae*) were also assayed individually and in pairwise combinations.

Firstly, yeast strains were streaked out on *Sabouraud* medium plates and grown at 28°C during 24h. After that, single colonies were used to inoculate precultures into 3 ml SGM medium on 12-well plates. After 24h of growth at 25ºC under orbital shaking (100 rpm), yeast cultures were combined, according to the community design, in 2 mL tubes. Then, 10 μL of the prepared mix culture (synthetic yeast consortia) were inoculated into 96-well plates to reach a final OD_600nm_ of 0.02 in 250 μL of SGM (each non-*Saccharomyces* strain member of the community was at a final OD_600nm_ of 0.004). Then, when necessary, 10 μL of the adjusted *S. cerevisiae* cultures were also added into the assays, together with the background community, to reach a final OD_600nm_ of 0.001 (that is, a 1:4 ratio with each non-*Saccharomyces* strains in the background, to simulate the lower abundance of *S. cerevisiae*, in relation to the rest of non-*Saccharomyces* yeasts found in natural yeast communities of grape musts). OD _600nm_ of the cultures were monitored during 168h as described previously for the growth phenotyping.

### Analysis of the function of synthetic yeast consortia

Sugar consumption was measured as the main ecological function of the assayed synthetic consortia, by the quantification of total fermentable sugars (glucose+fructose) consumed after 168h of fermentation (results shown in Figures 3 and 4). We used the DNS (3,5-dinitrosalicylic acid) method (Luyt et al., 2021), adapted to 96-well plates. Briefly, in each well 100 uL of the DNS solution (30 g/L potassium sodium tartrate; 16 g/L NaOH; 10 g/L DNS) were incubated (100ºC during 30 minutes) with 10 uL of each sample, then cooled at 4ºC for 30 minutes, and, finally, absorbance was measured at 540 nm in the microplate reader Varioskan Flash Multimode Reader (Thermo Scientific, USA). The absorbance values were interpolated in a calibration curve prepared using the same protocol. Ten half serial dilutions of standard SGM (200 g/L glucose+fructose) diluted with SGM without sugars by triplicate were used for the calibration curve. Concentration values above 12.5 g/L were out of the linearity range of the calibration curve. Therefore, all the assays exceeding this value were measured again, with a previous dilution of 1/10 or 1/20 as necessary. The fraction of sugars consumed by the assembled communities was calculated by the difference between the initial sugar concentration (200 g/L) and the final residual sugar concentration at the end of the fermentation assays, and normalised to 0-1 range (Table S8). The functional effect of adding one strain to a specific background community on the ecological function (fraction of sugars consumed) was expressed as dF_i_=F_j+i_ – F_j_ ; where F_j+i_ is the function of the background community j when the strain i is added and F_j_ is the function of the background community j.

### Predicting the function of complex consortia by a simple additive model

To predict the function of the assembled communities, we firstly created a simple model feed with the individual effect of each strain in co-cultures with *S. cerevisiae* (dF). Thus, as an example, the function of the community A (composed by the strains i, j and z, and co-inoculated with the *S. cerevisiae* strain) was predicted by this model as follows: F_A_=dF_i_ + dF_j_ + dF_z_ + F_Sc_. To test the accuracy of predictions based on this model, we calculated the Root Mean Square Error (RMSE) by using the rmse function of the *Metrics* package in R (v 4.2.2), that compares the observed values of the function of the communities and the prediction values using the additive model. High RMSE values indicate a low predictability accuracy of the model. For the individual predictions for each community we calculated the squared absolute difference between the observed and the predicted function value (represented against strain complexity in Figure 3E, Figure 4D and Figure S9).

### Estimation of Functional Effect Equations and function prediction

By following the methodology described by Diaz-Colunga et al. (2022), the functional effect of each strain (dF) is represented against the function of the different background communities where they are added F (A) (Figure 4A-B). Thus, the functional effect of the different strains (henceforth Functional Effect Equations, FEEs) can be defined mathematically as: dF_i_ (A)= a_i_ + b_i_ + F (A) + ϵ_i_ (A) where dF_i_ (A) is the effect of the strain i in the ecological function of the community A. The intercepts (a_i_) and the slope (b_i_) can define the ecological effect of each strain. ϵ_i_ (A) refers to the deviation of the residuals from the fit. As described by Diaz-Colunga et al. (2022), we tried this methodology to predict the function of the yeast communities assembled based on their taxonomic composition. Thus, by concatenating the FEEs of each strain that composes the community, we can estimate the ecological function (fraction of sugars consumed) of any randomly assembled community (composed by the 12 strains assayed). In order to double check the accuracy of this FEEs-based prediction model (Figure 4D), 131 new communities -not assembled in the first experiment-were assembled and assayed (Table S9). Following the methodology described above, the fraction of sugars consumed was calculated after 168h of SGM fermentation. To also test the replicability of the procedures, this new experiment was conducted by a different lab member. The predicted function values were then compared to the actual measured values to estimate the prediction ability of this model.

### Importance of phenotypic trait as predictors of FEEs slope and intercept

To assess the importance of all the individual phenotypic traits measured (those represented in Figure 2) as predictors of the functional effect of yeast strains on different ecological contexts (i.e. FEEs slope and intercept), we used random forest classification analysis using *randomForest* (v 4.7-1.1) package in R (Figure S12). We included the environmental preferences and the measured physico-chemical parameters (fermentation performance) as predictors of the slope and intercepts of the predicted function (sugar consumption), considered as the response variables. The growth efficiency values in the media used to test the growth ability using different organic acids and amino acids as carbon and nitrogen sources, were grouped, respectively, as carbon and nitrogen sources.

