## Supplementary Figures for "Predictability of the community-function landscape in wine yeast ecosystems"

### Supplementary Material

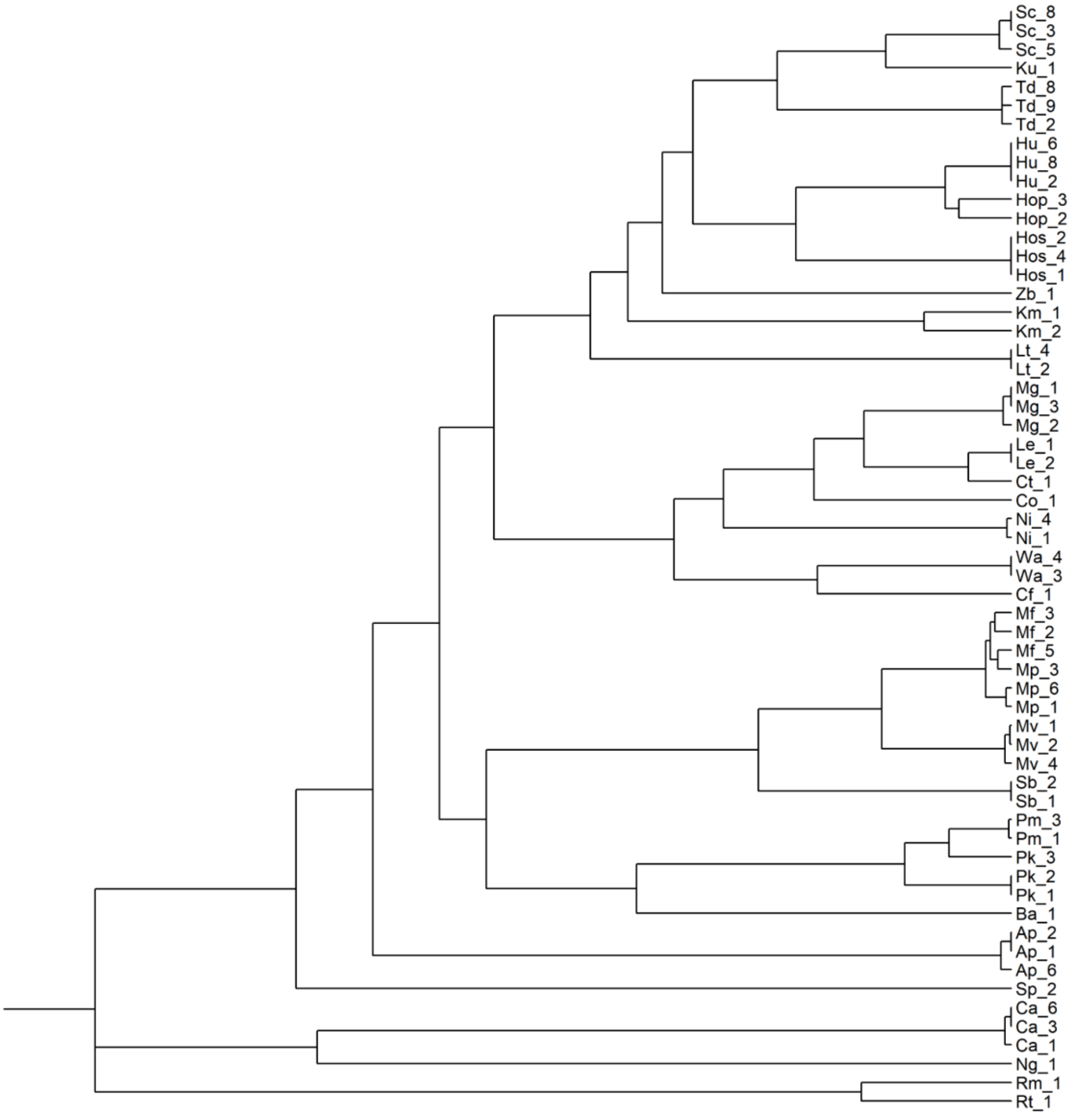


**Figure S1. Phylogenetic relationships representation of the 60 strains used in this work.** The phylogenetic tree was constructed using the 26s rRNA sequences of the 60 yeast strains (Table S1). The 60 sequences aligned were used to construct a maximum likelihood phylogenetic tree. Consensus tree was rooted using the Basidiomycota group as an outgroup.

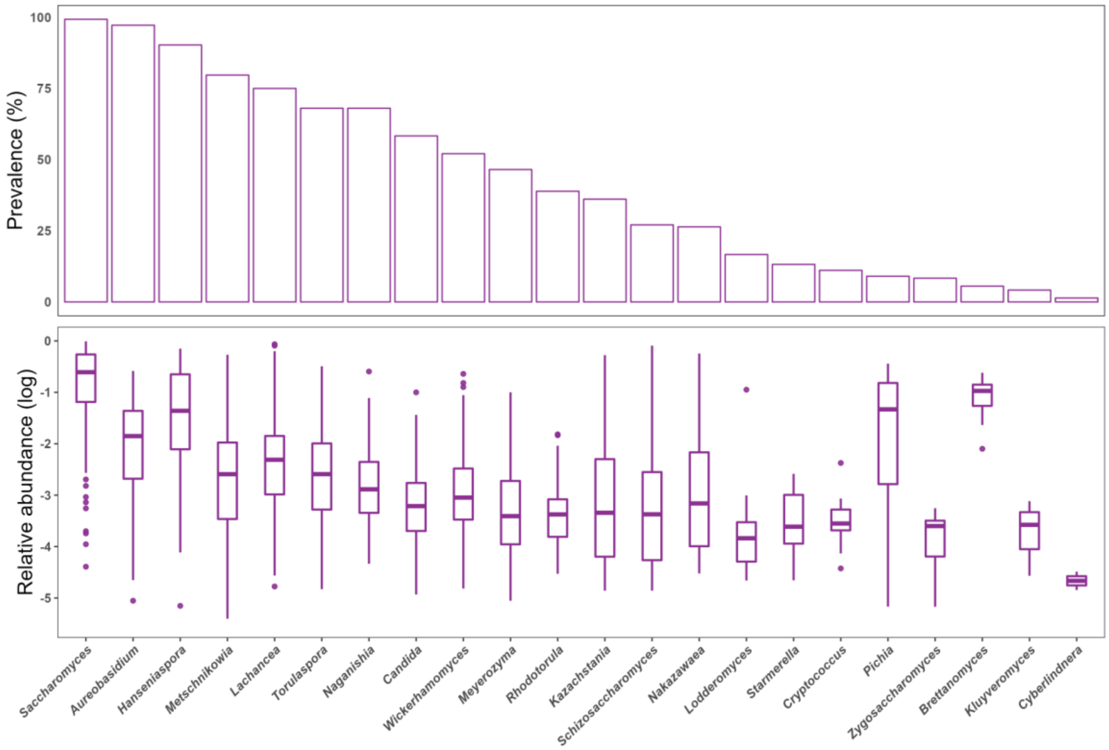


**Figure S2. All the strains included in the collection of this work belong to yeast genera that are found, to a greater or lesser extent, in wine alcoholic fermentation.** We show the estimated figures of population prevalence and relative abundance for the 22 yeast genera included in the collection, inferred using the ITS-amplicon data published by de Celis et al. (2022) from a large survey of 144 wine alcoholic fermentation samples. In the case of the genus *Metschnikowia*, data were obtained from Vicente et al., 2020.

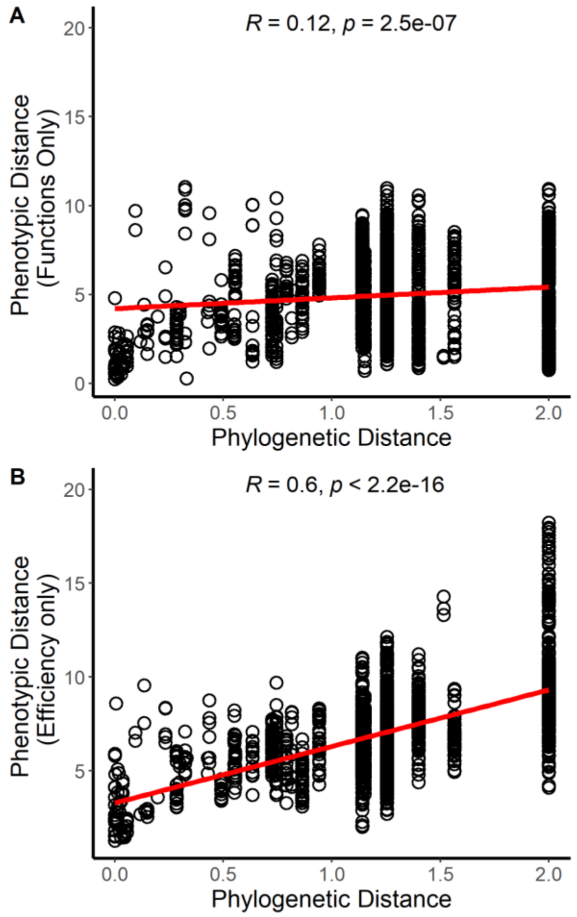


**Figure S3. Phylogenetic distance among the strains significantly correlated to their phenotypic distance**. Correlation between phylogenetic distance (the sum of branch lengths between strains pairs in the phylogenetic tree, Figure S1) and the phenotypic distance (Euclidean distance between the strains in the matrix of the phenotypic trait measured for each strain). (**A**) represents the phenotypic distance including just the wine physical-chemical parameters measured after 168h of growth in SGM (functions). (**B**) represents the phenotypic distance including just the efficiency growth value under the 28 environmental preferences assays. Dots represent each strain and red lines represent the Pearson correlation.

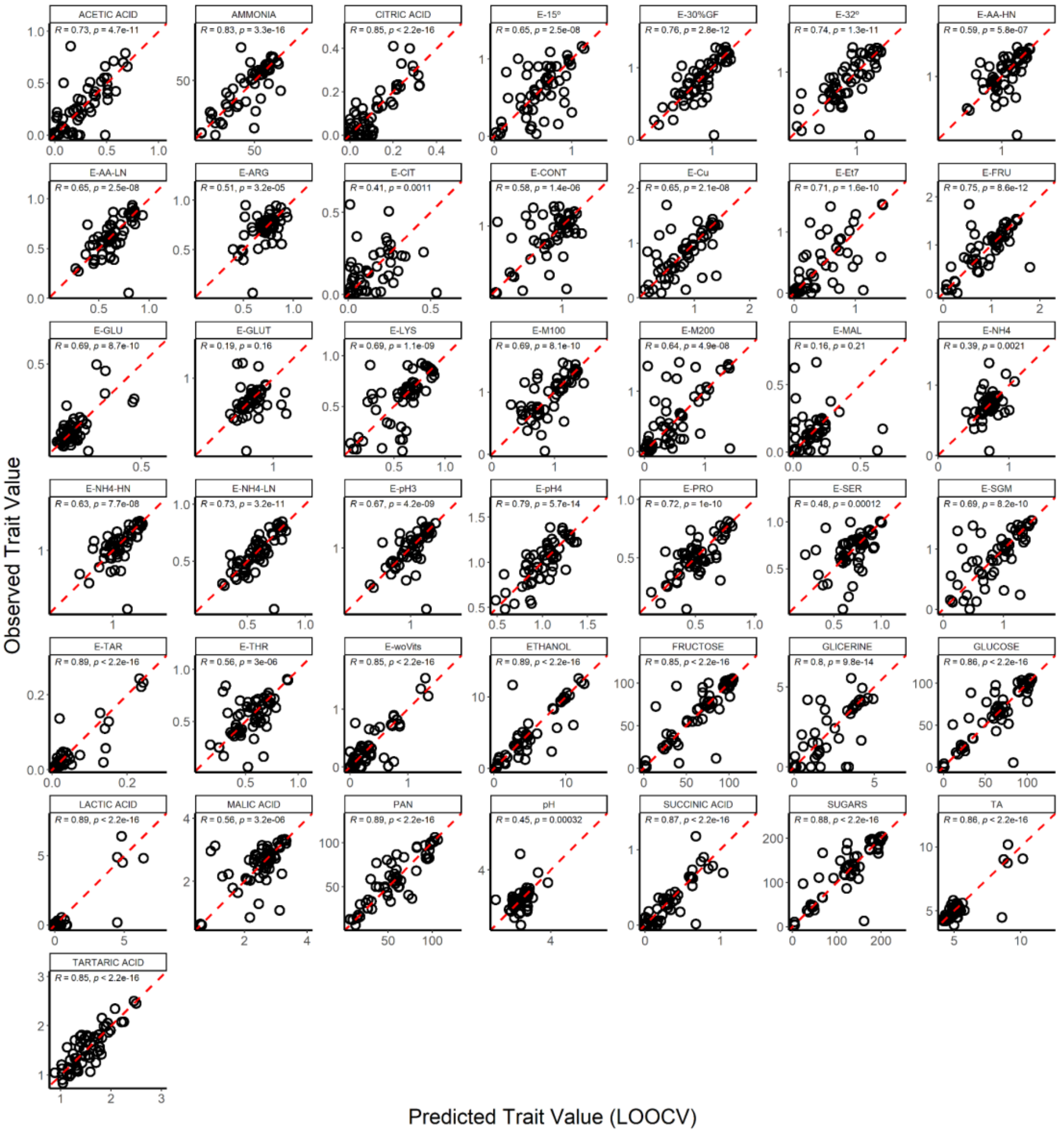


**Figure S4. Most of the phenotypic traits in the wine yeast collection of the study are predictable using the phylogeny.** Panel of predicted vs observed traits (28 environmental preferences analysed and 15 wine related parameters analysed after Synthetic Grape Must fermentation). Just the efficiency growth using glutamine (E-Glu) as sole nitrogen source and the efficiency growth using malic acid as sole carbon source (E-MAL) were not accurately predicted based on the phylogeny ($R$=0.19, $p$=0.16 and $R$=0.16, $p$=0.21, respectively). Predictions were made using phylogenetic imputation and tested by the leave one out-cross validation (LOOCV).

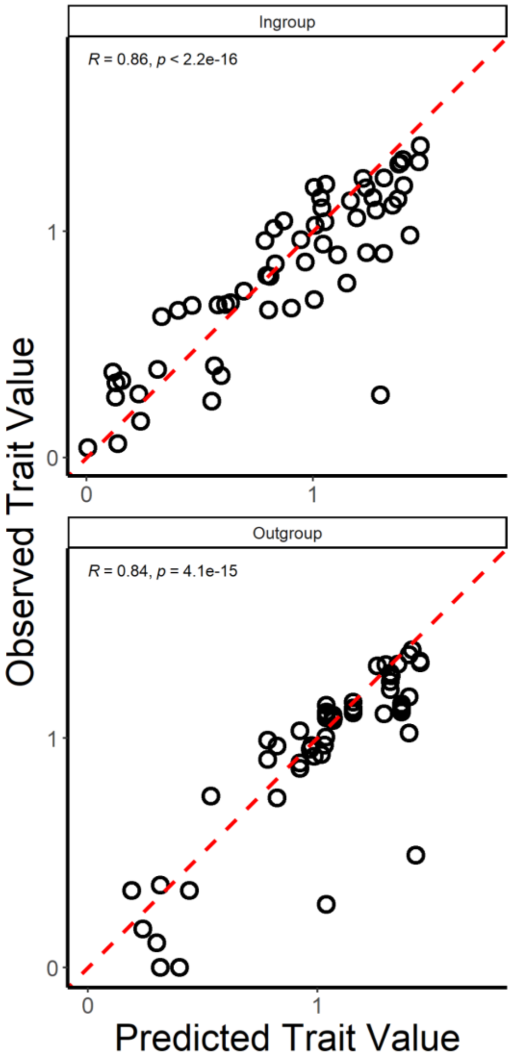


**Figure S5. The prediction based on phylogeny allows to predict the phenotype of new strains with a similar accuracy as replicate measures.** To check the accuracy of our phylogenetically-based prediction model, we assayed the growth efficiency in SGM of a wider collection of strains (the 60 strains already characterised and 53 new strains (Table S6)). In this way, we characterised this trait in 60 strains in two independent batches of experiments. The upper plot represents the prediction of this trait in the 60 strains based on the efficiency values of the second batch of the experiment. The lower plot represents the prediction of the 60 strains based on the phylogenetic relationships of the 53 new strains.

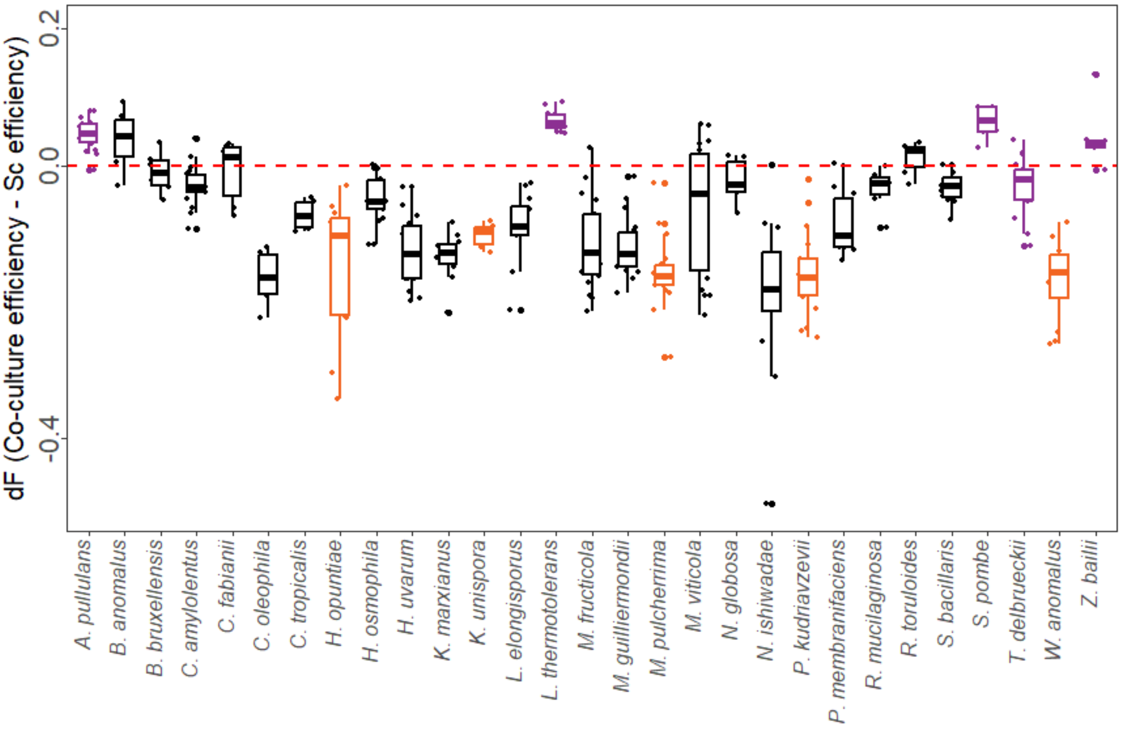


**Figure S6. Some wine yeast species cause a negative impact on the growth efficiency of *S. cerevisiae*.** To explore the effect of non-*Saccharomyces* strains on *S. cerevisiae* growth, we co-inoculated each non-*Saccharomyces* strain with *S. cerevisiae* Sc_5 strain. The functional effect (dF) of the different species were calculated by comparing efficiency values (total variation in cells density) of the co-cultures and the efficiency values of *S. cerevisiae* individual culture (Sc_5xSc_5), after 168h of SGM fermentation (Table S7). Purple and orange colours represent those species from which we selected strains with an enhanced or negative effect on *S. cerevisiae* co-culture performance, respectively. Boxplots represent the median and standard deviation of all the replicates of the different strains belonging to the same species.

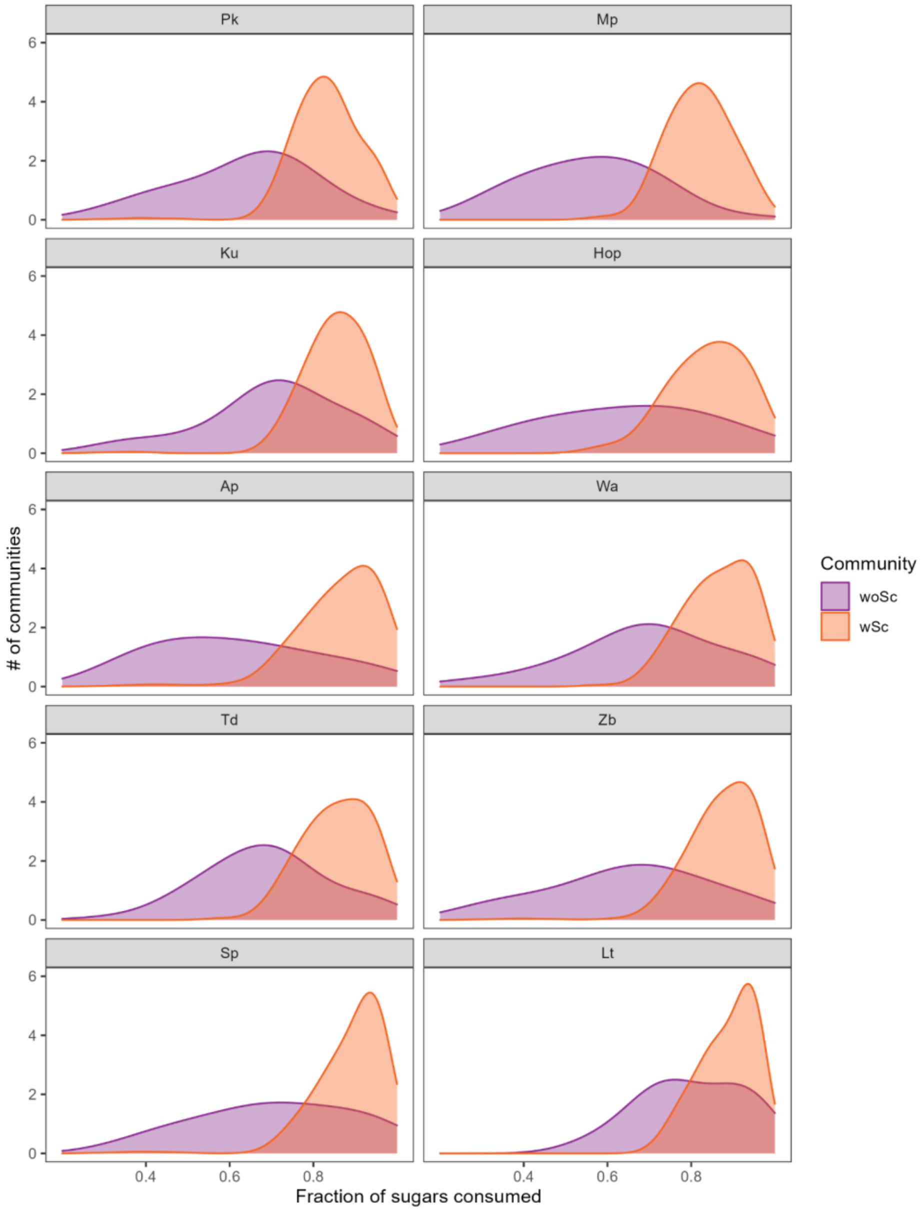


**Figure S7. Strains were not equally distributed among the communities based on their ecological function.** We found that some strains tend to be distributed in communities with lower function (less than the 70-60% of the sugars consumed after the fermentation) and other strains tend to be distributed in higher function communities (more than the 80-90% of the sugars consumed). This distribution is not explained by the individual function of strains (Figure 3D), therefore ecological interactions might emerge from complex communities.

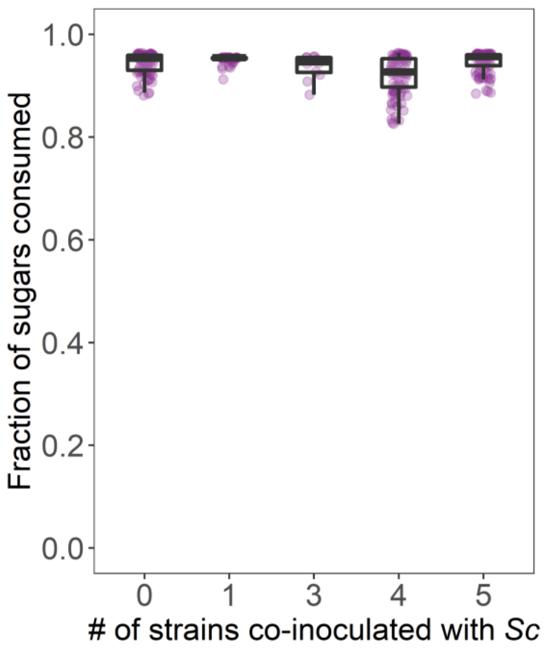


**Figure S8. Community diversity does not affect function in communities that harbours neutral-effect strains.** We just represented the communities that do not contain Hop, Mp, Pk strains (strains that cause a negative effect on *S. cerevisiae* co-culture function, Figure 3D). Each dot represents the fraction of sugars consumed by each community in which *S. cerevisiae* strains were inoculated.

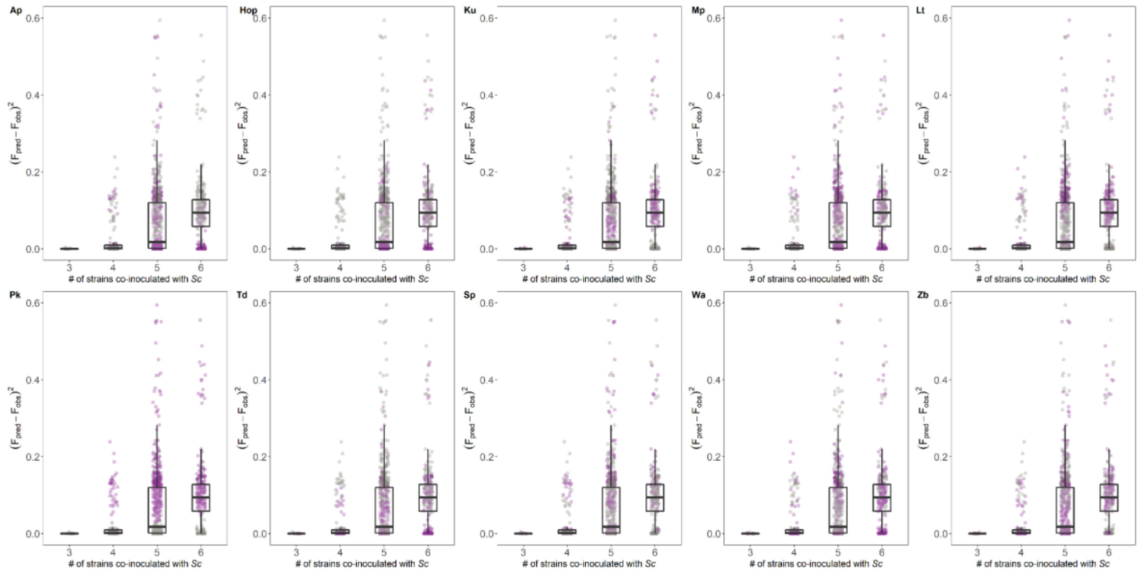


**Figure S9. Contribution of the different non-*Saccharomyces* strains to the prediction accuracy of the additive model at the different levels of complexity of the communities inoculated with *S. cerevisiae*.** We can observe how the different strains are distributed among the communities whose function (fraction of sugars consumed) is predicted better (low $F_{pred}$-$F_{obs}$ values) or worse (high $F_{pred}$-$F_{obs}$ values). We can observe that the presence of *Pk* causes a marked loss on how predictable the community is. The strong pattern of this strain can be explained by the fact that the strong detrimental effect of the strain *Pk* on both *S. cerevisiae* strain co-culture (Figure 4D) is diluted under complex community context as a consequence of the emergence of higher-order interactions.

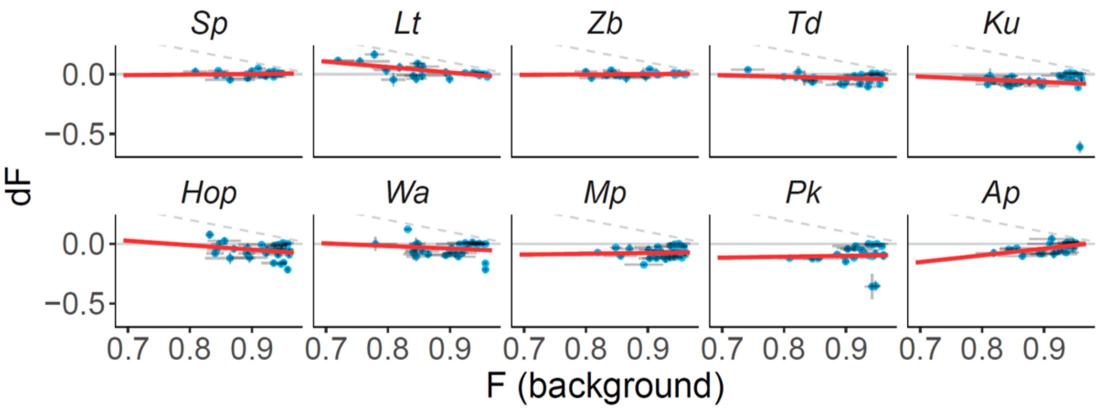


**Figure S10. Functional Effect Equations of the strains on yeast communities that contain *S. cerevisiae*.** When we explore the FEEs of the strains just considering communities that contain *S. cerevisiae*, the clear patterns of the strains observed in Figure 4B are lost (as *S. cerevisiae* is the main contributor to the ecological function of wine fermentation). However, some strains (*Hop*, *Mp*, *Pk* and *Ap*) still show marked negative patterns in the function of the communities regardless of the function of the background community.

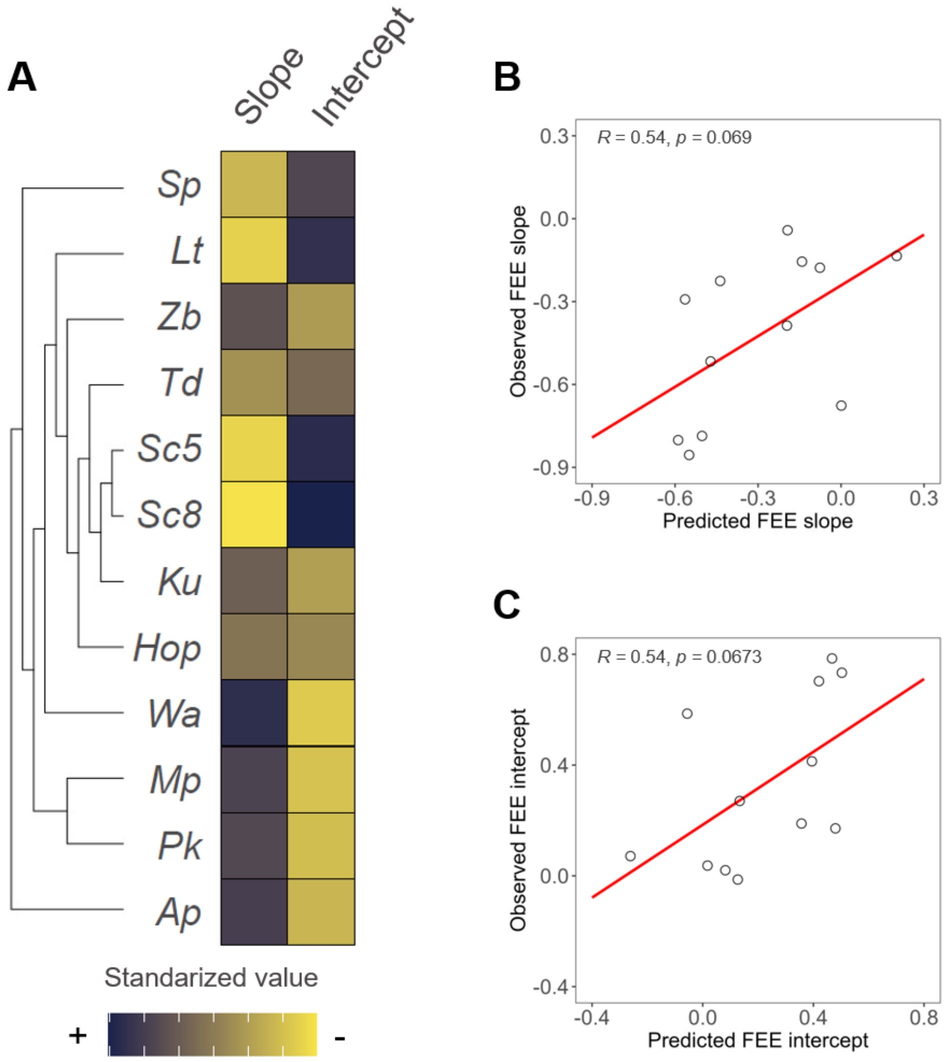


**Figure S11. Functional effects of the strains on their ecological context (FEEs slopes and intercepts) are less predictable from phylogeny than the individual phenotypic traits.** Although correlation signal is observed between the observed and the predicted based on phylogeny FEEs parameters, the functional effect of strains cannot be clearly predicted using the phylogeny imputation. The strong impact of the function of the background communities might explain this loss of predictability when we consider the ecological effect of the strains.

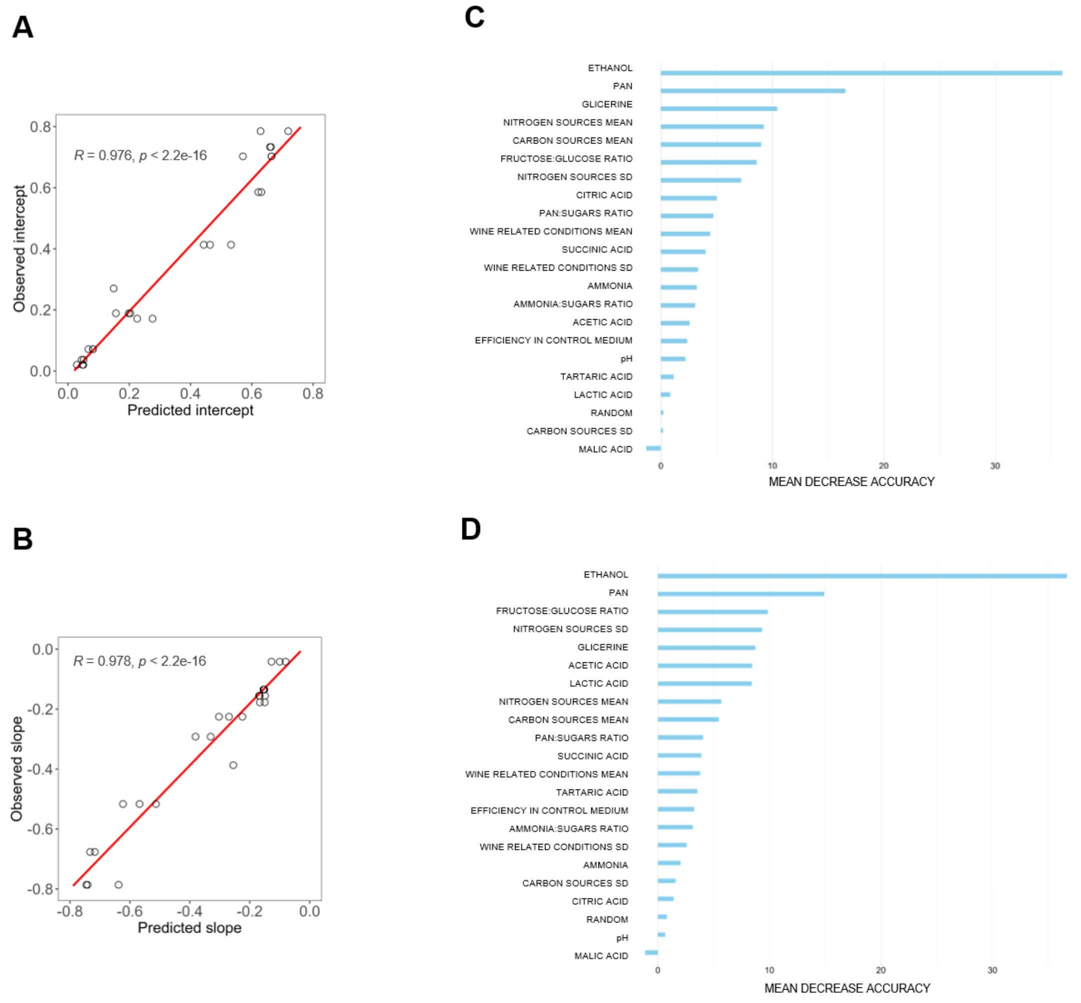


**Figure S12. Ethanol production and Primary Amino acid Nitrogen consumption are the phenotypic traits that better explain the ecological effect of the strains in wine yeast communities.** A random forest model was used to predict the FEEs parameters; intercept **(A)** and slope **(B)** based on the 43 phenotypic traits analysed in this study (Figure 2A-B). The percentage of model accuracy that decreases when we leave out each trait is represented for the intercept prediction **(C)** and for the slope prediction **(D)**.

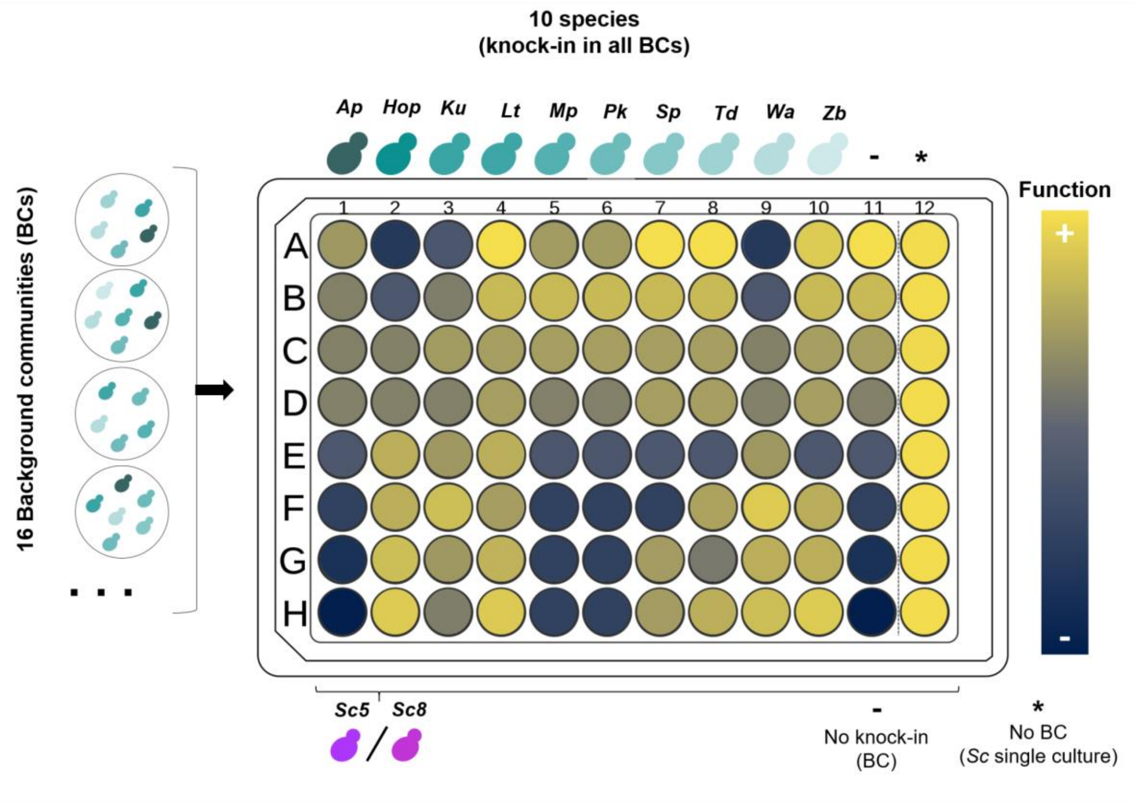


**Figure S13. Experimental design for the study of ecological effect of strains on different yeast background communities.** A total of 16 different background communities (composed with 3, 4, 5 or 6 non-*Saccharomyces* strains) were inoculated in Synthetic Grape Must on 96-well plates. In addition, every single strain was co-inoculated in the wells containing each background community, in order to increase the number of possible combinations of strains. Three plates with this display were prepared; one plate was inoculated with Sc5 *S. cerevisiae* strain, another plate was inoculated with Sc8 *S. cerevisiae* strain, and another plate was not inoculated with any *S. cerevisiae* strain. Thus, we created a total of 528 different communities, inoculated by triplicate. After 168h of fermentation, the ecological function of each community was calculated by measuring the residual sugars (glucose+fructose). Yellow colour indicates high function communities (a high fraction of sugars was consumed after fermentation), blue colour indicates low function communities (a low fraction of sugars was consumed after fermentation). The functions of all the communities, pairwise and individual assays are shown in Table S8.
